# Comparative analysis reveals the species-specific genetic determinants of ACE2 required for SARS-CoV-2 entry

**DOI:** 10.1101/2020.09.20.297242

**Authors:** Wenlin Ren, Gaowei Hu, Xiaomin Zhao, Yuyan Wang, Hongyang Shi, Jun Lan, Yunkai Zhu, Jianping Wu, Devin J. Kenney, Douam Florian, Yimin Tong, Jin Zhong, Youhua Xie, Xinquan Wang, Zhenghong Yuan, Dongming Zhou, Rong Zhang, Qiang Ding

**Author notes:** Corresponding authors (Q.D.); (R.Z.). These authors contributed equally to this work (Wenlin Ren, Gaowei Hu, Xiaomin Zhao).

## Abstract

Coronavirus interaction with its viral receptor is a primary genetic determinant of host range and tissue tropism. SARS-CoV-2 utilizes ACE2 as the receptor to enter host cell in a species-specific manner. We and others have previously shown that ACE2 orthologs from New World monkey, koala and mouse cannot interact with SARS-CoV-2 to mediate viral entry, and this defect can be restored by humanization of the restrictive residues in New World monkey ACE2. To better understand the genetic determinants behind the ability of ACE2 orthologs to support viral entry, we compared koala and mouse ACE2 sequences with that of human and identified the key residues in koala and mouse ACE2 that restrict viral receptor activity. Humanization of these critical residues rendered both koala and mouse ACE2 capable of binding the spike protein and facilitating viral entry. The single mutation that allowed for mouse ACE2 to serve as a viral receptor provides a potential avenue for the development of SARS-CoV-2 mouse model.

## INTRODUCTION

Coronaviruses, belonging to the *Orthocoronavirinae* subfamily and *Coronaviridae* family^1,2^, are enveloped viruses with an approximately 30 kb, positive-sense, single-stranded RNA genome^3,4^. In the last two decades, three coronaviruses— severe acute respiratory syndrome coronavirus (SARS-CoV), Middle East respiratory syndrome coronavirus (MERS-CoV) and SARS coronavirus 2 (SARS-CoV-2)—have caused outbreaks of severe diseases in humans^5-8^. SARS-CoV-2, responsible for the ongoing COVID-19 pandemic, poses a continuing threat to global public health, especially since there are no specific and effective clinical therapeutics or approved vaccines. To facilitate drug and vaccine development, there is an urgent need to create novel models for studying the basic biology and pathogenesis of SARS-CoV-2^9,10^.

The first and essential step of virus infection is cellular receptor recognition. It has been demonstrated that the interaction of a virus with (a) species-specific receptor(s) is a primary genetic determinant of host tropism and therefore constitutes a major interspecies barrier at the level of viral entry^11-13^. SARS-CoV-2 enters host cells by binding angiotensin-converting enzyme 2 (ACE2) in a species-dependent manner^6,14,15^. Murine ACE2 does not efficiently bind the SARS-CoV-2 spike (S) protein, hindering viral entry into murine cells; consequently, human ACE2 transgenic and knock-in mice have been developed to study *in vivo* the infection and pathogenesis of SARS-CoV-2^16-19^.

In exploring the host tropism of SARS-CoV-2, we and others have reported that a diverse array of ACE2 orthologs can serve as a receptor to mediate SARS-CoV-2 entry, suggesting a broad host range^20-22^. We showed that ACE2 orthologs from New World monkeys (marmoset, tufted capuchin and squirrel monkey) or koalas cannot support infection with SARS-CoV-2 due to incompatibility of the S protein to the orthologs^20^. This explains, at least in part, the observation that marmosets are resistant to experimental SARS-CoV-2 infection^23^. We identified the restrictive residues at positions 41 and 42 in New World monkey ACE2 orthologs, which disrupt the formation of hydrogen bonds observed between human ACE2 and SARS-CoV-2 S protein. Replacing these two residues with the corresponding human counterparts resulted in a gain-of-function phenotype that permitted these ACE2 orthologs to act as functional SARS-CoV-2 receptors, as evidenced by binding of the viral S protein and subsequent viral entry ^20^. Such findings could inform the development of marmosets as a SARS-CoV-2 infection model. Interestingly, the molecular basis for the inability of koala and mouse ACE2 to support viral entry is different from that of New World monkeys. In this study, we aimed to identify the determinants responsible for the restricted activity of koala and mouse ACE2 as a SARS-CoV-2 receptor by genetic and functional analysis. Our work could provide greater insight into the species-specific restrictions of SARS-CoV-2 cell entry and also inform animal model development.

## RESULTS

### Comparative analysis identifies the potential species-specific residues in koala and mouse ACE2 that restrict SARS-CoV-2 entry

New World monkey, koala, and mouse ACE2 cannot function as a SARS-CoV-2 receptor to mediate entry, but New World monkey ACE2 humanized at residues 41 and 42 could support viral entry^20^. To identify the genetic determinants that restrict the viral receptor activity of koala and mouse ACE2, we analyzed ACE2 orthologs (**Fig 1a**), especially the residues that interface with the SARS-CoV-2 receptor-binding domain (RBD) of the spike protein, across a diverse set of species (**Fig 1b and Supplemental Fig 1**). Based on the structure of human ACE2 complexed with the SARS-CoV-2 RBD^24-27^, K31 and K353 of ACE2 are located in the contact interface with the RBD and provide a substantial amount of energy to stabilize the ACE2-spike complex^28^. Specifically, K31 and K353 form hydrogen bonds with Q493 and the carboxyl oxygen of G502 of the SARS-CoV-2 RBD, respectively. ACE2 Y83 also contributes to the interaction by forming hydrogen bonds with SARS-CoV-2 RBD N487 and Y489 as well as a π-π interaction with F486 (**Fig 1c**). However, some of these key residues differ in koala and mouse ACE2: T31 and F83 in koala, and F83 and H353 in mouse. It is conceivable that these substitutions could disrupt the interaction of these ACE2 orthologs with the viral spike and thus impair viral entry.

**Figure 1.**
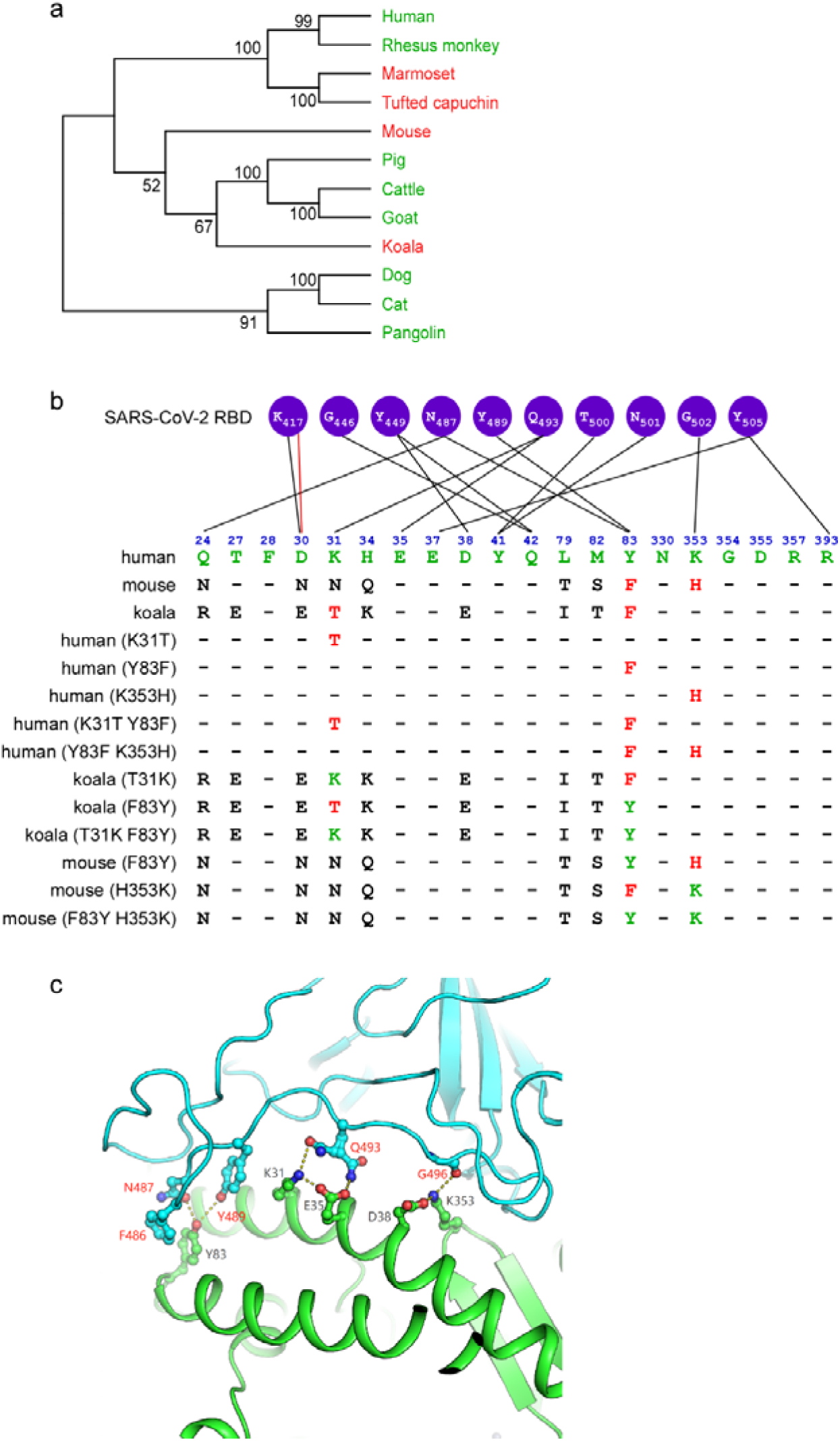
The potential residues in ACE2 that restrict SARS-CoV-2 entry. (a) A phylogenetic tree was constructed based on the protein sequences of ACE2 orthologs by using the Neighbor-joining method conducted in program MEGA7^43^. The percentage of replicate trees in which the associated taxa clustered together in the bootstrap test (1000 replicates) are shown next to the branches. The ACE2 sequences of these species were downloaded from NCBI, and accession numbers are shown in Fig. S1. (b) Alignment of the residues of human, koala and mouse ACE2 at the interface of ACE2 with the SARS-CoV-2 spike protein (first three rows). The restrictive residues of koala or mouse ACE2 are highlighted in red. The favorable residues of human ACE2 are highlighted in green. A series of mutant ACE2 orthologs bearing restrictive or favorable residues were constructed in this study (remaining rows). (c) Cartoon of the binding interface between human ACE2 and the SARS-CoV-2 receptor-binding domain (RBD) (PDB code: 6M0J). ACE2 and the SARS-CoV-2 RBD colored in green and cyan, respectively. Key residues (K31, Y83, and K353) discussed in this study and their interacting residues are shown as ball-and-stick representations.

### Humanization of restrictive residues in ACE2 orthologs rescue binding to the SARS-CoV-2 and SARS-CoV spike protein

To directly uncover the impact of these differing residues in koala and mouse ACE2 on viral entry, we introduced these residues into human ACE2, generating the following mutants: hACE2 (K31T), hACE2 (Y83F), hACE2 (K353H), hACE2 (K31T/Y83F) hACE2 (Y83F/K353H) (**Fig 1b**). Conversely, we generated mutant koala and mouse ACE2s bearing the corresponding human residues: kACE2 (T31K), kACE2 (F83Y), kACE2 (T31K/F83Y); mACE2 (F83Y), mACE2 (H353K), mACE2 (F83Y/H353K). We then tested the ability of these mutants to bind SARS-CoV-2 S protein and mediate virus entry.

As binding of SARS-CoV-2 S protein to human ACE2 is a prerequisite for viral entry^6,14^, we employed a cell-based assay using flow cytometry to assess the binding of S protein to the different ACE2 orthologs and mutants. We cloned the cDNA of our ACE2 sequences, each with a C-terminal FLAG tag, into a bicistronic lentiviral vector that expresses the fluorescent protein zsGreen1 driven by an IRES element (pLVX-IRES-zsGreen1) to monitor transduction efficiency. Next, a purified fusion protein consisting of the S1 domain of the SARS-CoV-2 S protein and an Fc domain of human lgG (S1-Fc) was incubated with A549 cells transduced with the various ACE2 lentiviruses after 48 hours transduction (**Fig 2a**). Binding of S1-Fc to ACE2 was then determined by flow cytometry as the percent of cells positive for S1-Fc among the ACE2-expressing cells (zsGreen1+ cells) (**Fig 2b**). Additionally, by immunoblotting, we confirmed that the expression of each FLAG-tagged ACE2 ortholog and variant was comparable in the transduced cells (**Fig 2 c, d**). Thus, any differences in SARS-CoV-2 cell entry activity we observed were not simply due to differences in protein expression.

**Figure 2.**
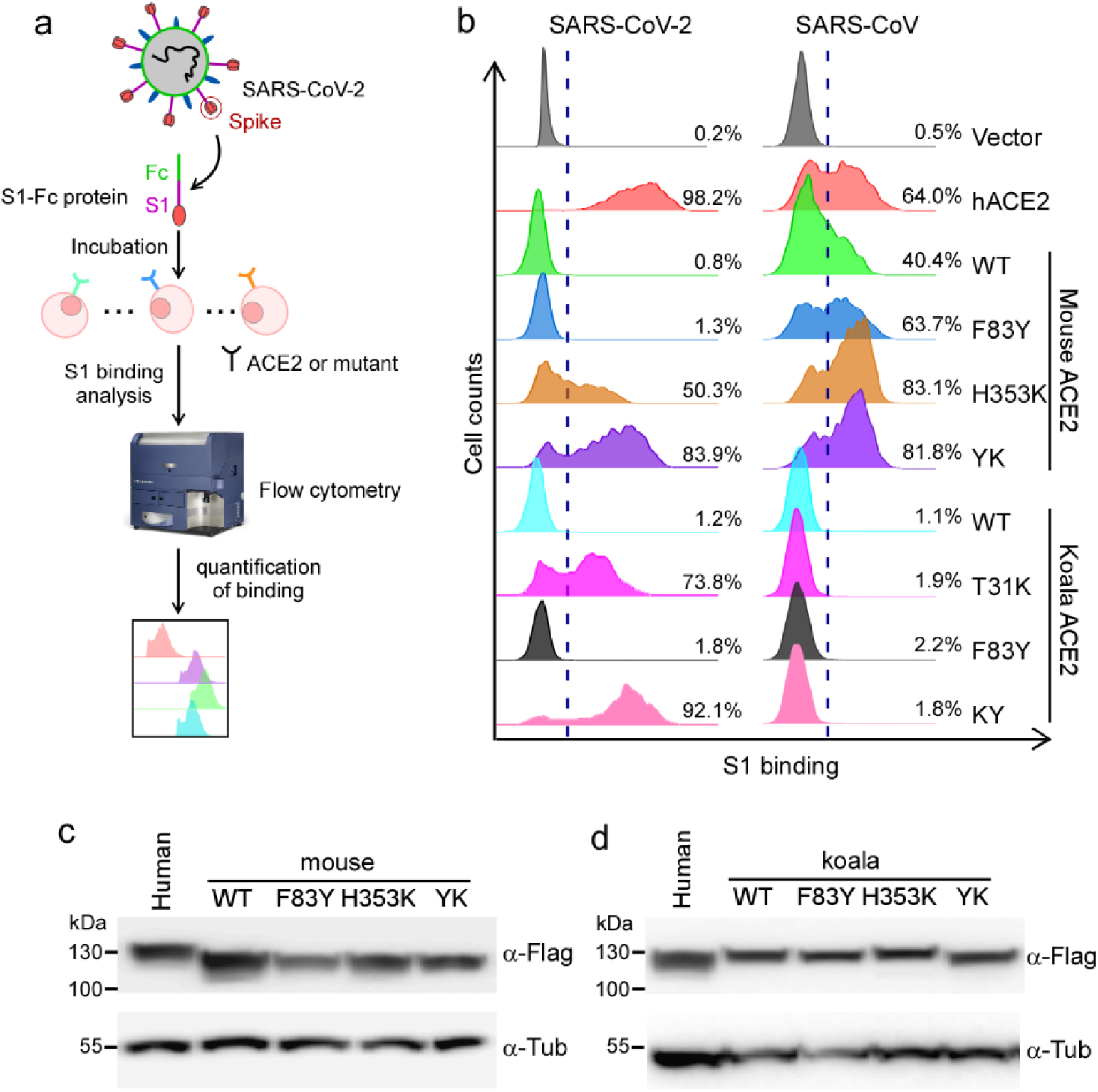
ACE2 mutants bind viral spike protein. (a-b) A549 cells were transduced with ACE2 orthologs or their mutants as indicated, incubated with the recombinant S1 domain of the SARS-CoV-2 or SARS-CoV spike protein C-terminally fused with Fc, and then stained with goat anti-human IgG (H + L) conjugated to Alexa Fluor 647 for flow cytometry analysis. Values are expressed as the percent of cells positive for S1-Fc among the ACE2-expressing cells (zsGreen1+ cells). (c-d) Representative immunoblots of A549 cells transduced with lentiviruses expressing FLAG-tagged ACE2 orthologs and humanized mutants were subjected to immunoblotting. Tubulin served as the loading control.

Consistent with our previous results^20^, the binding of S1-Fc to A549 cells expressing mouse or koala ACE2 was very low and comparable to levels observed in cells transduced with empty vector, whereas S1-Fc protein efficiently bound to A549 cells expressing human ACE2 (**Fig 2b and Supplemental Fig 2**). As shown in Supplemental Fig 2, mutating human ACE2 at position 83 to the corresponding residue in mouse or koala ACE2 (Y83F) did not dramatically affect binding with SARS-CoV-2 spike (98.5% vs 98.1%) whereas substitution to the mouse residue at position 353 (K353H) slightly decreased binding (75.1% vs 98.1%). Of note, the human-to-mouse hACE2 Y83F/K353H double mutant significantly decreased binding (49.9% vs 98.1%). The human-to-koala single hACE2 K31T mutant had dramatically impaired binding (52.8% vs 98.1%) that was further reduced with mutation of residue 353 (K31T/K353H; 28.6% vs 98.1%), indicating that K31 is critical for the ACE2-spike interaction.

To gain further insights into the genetic determinants of ACE2 as the SARS-CoV-2 receptor, we examined the binding of our humanized mouse or koala ACE2 mutants with the spike protein (**Fig 2a**). The humanized mouse or koala ACE2 mutated at residue 83 (mACE2 F83Y, kACE2 F83Y) did not show increased binding efficiencies compared to wild-type mouse or koala ACE2 (1.3% vs 0.8%; 1.8% vs 1.2%, respectively) (**Fig 2b, left panel**). However, the other singly humanized mutants (mACE2 H353K and kACE2 T31K) exhibited significantly improved binding with SARS-CoV-2 spike (50.3% and 73.8%, respectively). Notably, the doubly humanized mouse (mACE2 F83Y/H353) and koala (kACE2 T31K/F83Y) mutants demonstrated even greater binding efficiencies (83.9% and 92.1%). Together, these results indicate that although the F83Y mutation by itself had minimal effect on the binding of mouse or koala ACE2 to SARS-CoV-2 spike, combining this mutation with H353K or T31K, respectively, had a synergistic effect, greatly enhancing interaction with the viral spike to levels approaching that of human ACE2. Thus, we identified residue 353 in mouse ACE2 and residue 31 in koala ACE2 as the genetic basis for the species-specific restriction of these two orthologs in binding the SARS-CoV-2 spike.

To determine whether these observations were specific to the SARS-CoV-2 spike, we also assessed the abilities of these mutants to bind the SARS-CoV spike protein. In contrast to SARS-CoV-2 spike, mouse ACE2 could bind SARS-CoV spike (40.4% vs 0.8%) (**Fig 2b, right panel**). Single or double mutations (F83Y, H353K or F83Y/H353K) in mouse ACE2 further increased binding efficiency (63.7%, 83.1% or 81.8%, respectively), at times even exceeding that of human ACE2 (64.0%) (**Fig 2b, right panel**). This is in line with previous reports that humanizing four residues in rat ACE2 (82-84 and 353) converted this ortholog to an efficient SARS-CoV receptor ^29,30^. Intriguingly, unlike the SARS-CoV-2 spike, WT or humanized koala ACE2 displayed negligible binding to the SARS-CoV spike (**Fig 2b, right panel**). These observations imply that although SARS-CoV and SARS-CoV-2 both share human ACE2 as the cellular receptor for entry, there are functionally important differences in receptor interaction and recognition, which could contribute to differences in the host range, tissue tropism and pathogenesis of these two related viruses.

### Genetic modification of koala and mouse ACE2 renders cells susceptible to SARS-CoV-2 pseudovirus

Next, we sought to test whether ectopic expression of our ACE2 variants were functional and could promote entry of MLV retroviral particles (Fluc as the reporter) pseudotyped with the SARS-CoV-2 spike protein into A549 cells. For comparison, we also included VSV-G and SARS-CoV spike pseudotyped virons. A549 cells expressing the ACE2 variants were inoculated with the different pseudoparticles. At 48h after inoculation, the cells were lysed and the luciferase activity assayed to determine pseudovirus entry. As expected, VSVGpp readily infected all cells with similar efficiencies independent of which ACE2 ortholog or variant was expressed (**Fig 3**, dark gray columns). Compared to vector-transduced A549 cells, expression of human ACE2, but not mouse or koala, enhanced the entry of SARS-CoV-2 pseudoparticles (**Fig 3a and b**, black columns). These data were in line with the binding efficiencies of these ACE2 proteins we observed with the SARS-CoV-2 spike (**Fig 2b**). Neither the mouse nor the koala F83Y ACE2 variant significantly increased viral entry compared to the WT ortholog; in contrast, mACE2 H353K and kACE2 T31K each significantly enhanced viral entry, and the increased entry was maintained in the doubly humanized mutants (**Fig 3 a and b**).

**Figure 3.**
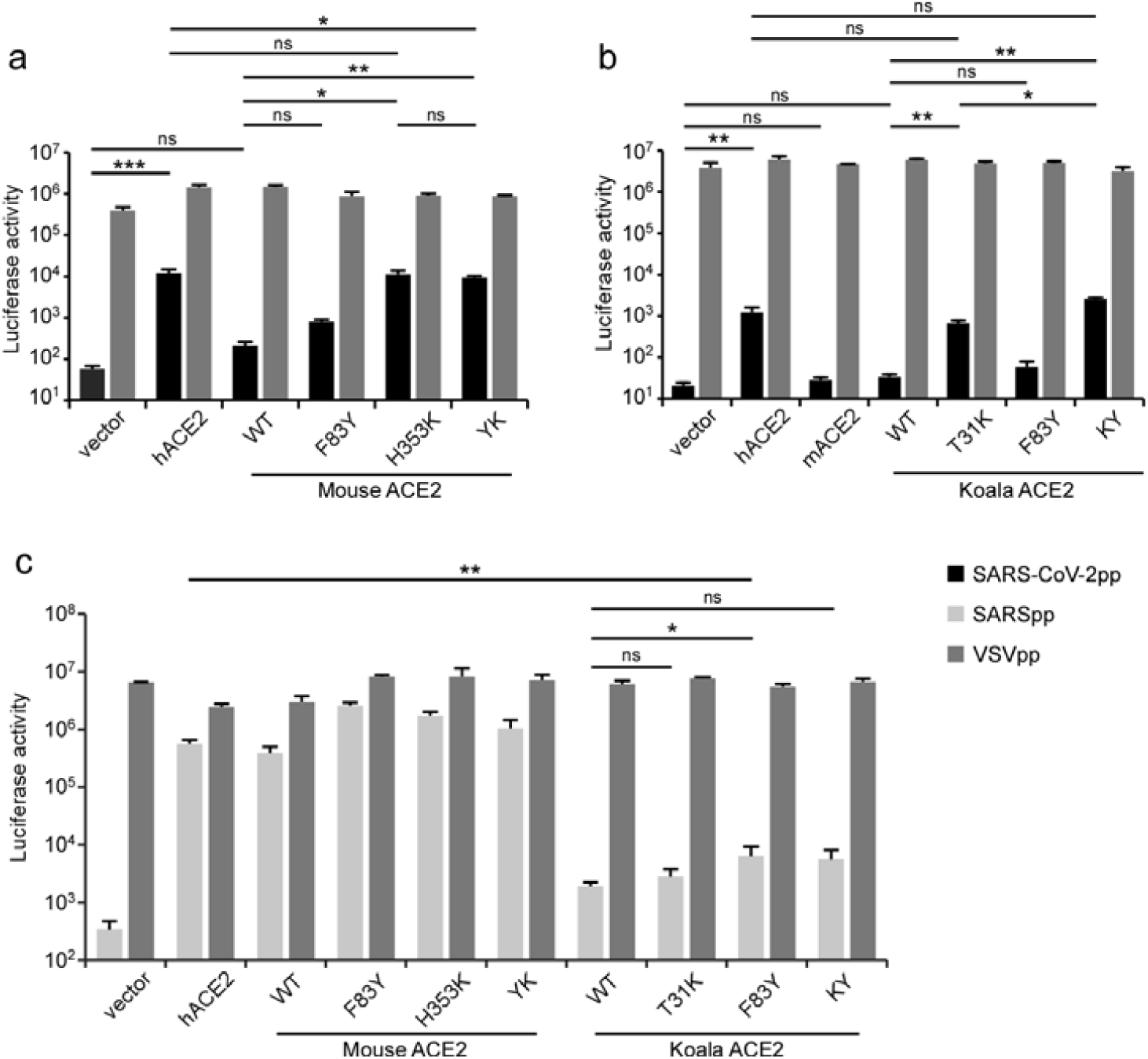
Ability of ACE2 orthologs and their humanized mutants to mediate entry of SARS-CoV-2 and SARS-CoV pseudoparticles. A549 cells transduced with mouse ACE2, koala ACE2 or their humanized mutants were infected with SARS-CoV-2 (a-b) or SARS-CoV (c) pseudoparticles. Two days after infection, cells were lysed and luciferase activity determined. VSV pseudoparticles were used as the control. All infections were performed in triplicate, and the data are representative of two independent experiments (mean ± standard deviation). ns, no significance; *, P < 0.05, **, P < 0.01, ***, P < 0.001. Significance assessed by one-way ANOVA.

As shown above (**Fig 2b**), SARS-CoV spike could bind mouse, but not koala, ACE2, and humanization of koala ACE2 at position 31 and/or 83 did not enhance binding with the SARS-CoV spike, unlike SARS-CoV-2. To further validate this observation, we tested the capabilities of ACE2 orthologs and their humanized versions to mediate SARS-CoV pseudoparticle entry. Consistent with the spike binding data, mouse ACE2 and its humanized mutants could mediate SARS-CoV pseudoparticle entry; however, the koala ACE2 mediated SARS-CoV entry with very low efficiency that was not enhanced with the T31K and/or F83Y variants (**Fig 3c**, light gray bars).

Collectively, the function of our ACE2 orthologs and their humanized mutants to mediate SARS-CoV-2 or SARS-CoV pseudoparticle entry corresponded with their ability to bind the respective viral spike proteins. Humanization of mouse or koala ACE2 by alteration of one or two amino acids resulted in the ability to mediate SARS-CoV-2 entry. Interestingly, mouse, but not koala, ACE2 could bind the SARS-CoV and SARS-CoV-2 spike proteins and mediate entry of pseudotyped particles for each. In contrast, humanization of koala ACE2 could bind only the SARs-CoV-2 spike protein and mediate entry of only particles pseudotyped with this protein. These observations highlight the different mechanisms used by SARS-CoV and SARS-CoV-2 for entry via ACE2.

### Humanization of restrictive residues in koala and mouse ACE2 orthologs renders cells susceptible to authentic SARS-CoV-2 infection

To directly test the ability of ACE2 orthologs to mediate SARS-CoV-2 entry during viral infection, we performed a genetic complementation experiment in A549 cells that lack endogenous ACE2 expression and are not susceptible to SARS-CoV-2 infection^31^.

A549 cells ectopically expressing our individual ACE2 orthologs or mutants were infected with SARS-CoV-2 (MOI=1). At 48 h post-infection, the complemented A549 cells were fixed and underwent immunofluorescent staining for intracellular viral nucleocapsid protein, an indicator of virus replication (**Fig 4a**). As expected, A549 cells expressing mouse or koala ACE2 were not susceptible to SARS-CoV-2 infection while those expressing human ACE2 were susceptible (**Fig 4b**). Consistent with the data from our binding and SARS-CoV-2 pseudoparticle experiments, A549 cells expressing humanized koala (T31K) or mouse (H353K) ACE2 were readily susceptible to SARS-CoV-2 infection; as expected, the F83Y mutation in koala (T31K) or mouse (H353K) ACE2 further enhanced infection efficiency. However, the F83Y single mutation in mouse or koala ACE2 had limited function (**Fig 4b and c**). Collectively, our results demonstrate that mouse and koala ACE2 with genetic humanization in the identified restrictive residues confer susceptibility to authentic SARS-CoV-2 infection.

**Figure 4.**
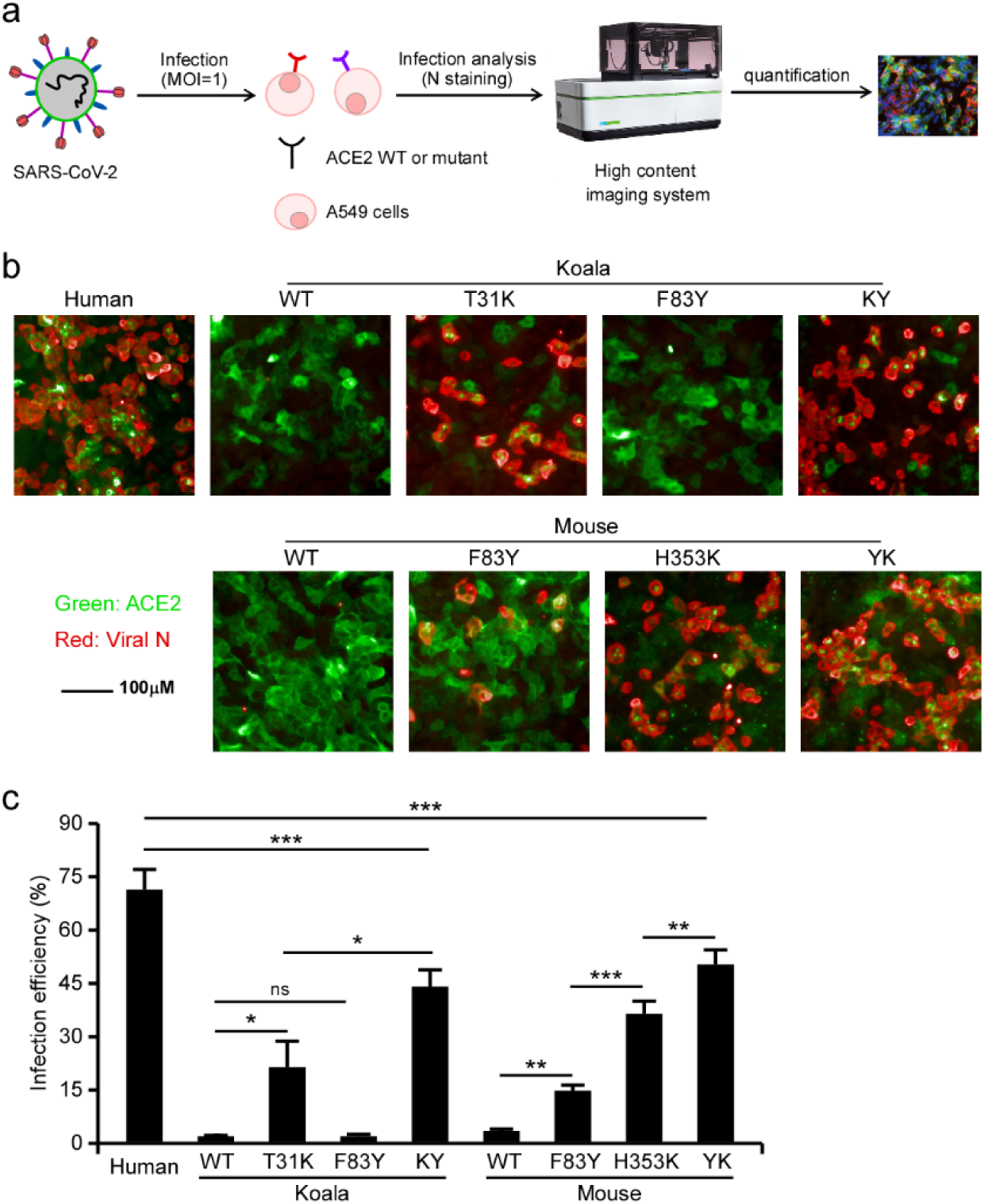
The capability of ACE2 mutants to facilitate viral entry. (a-b) A549 cells transduced with lentiviruses expressing ACE2 orthologs or humanized mutants were infected with SARS-CoV-2 virus (MOI=1). Expression of the viral nucleocapsid protein was visualized by immunofluorescence microscopy. Viral nucleocapsid (N) (red), and ACE2 (green) are shown. (c) The infection was quantified by a high-content imaging system. The graph shows the mean and SD (mean ± standard deviation) from two independent experiments performed in triplicate. ns, no significance; *, P < 0.05, **, P < 0.01, ***, P < 0.001. Significance assessed by one-way ANOVA.

## DISCUSSION

The host range of coronavirus is primarily determined by its interaction with cellular receptors^3,11,12,32,33^. SARS-CoV-2 utilizes ACE2 as the receptor to enter host cells, and it has been shown that a wide range of ACE2 orthologs from mammals can bind viral spike protein to facilitate entry^20-22,34^. However, New World monkey, koala, and mouse ACE2 exhibit limited support for SARS-CoV-2 entry^20^, and the underlying molecular mechanisms governing such restrictions are poorly understood. In this study, we conducted a systematic analysis of the genetic determinants of ACE2 that restrict the usage of koala and mouse ACE2 by SARS-CoV-2 for entry. We found that the genetic determinants responsible for the inability of koala and mouse ACE2 to support SARS-CoV-2 entry differ from those of New World monkeys (residues 41 and 42)^20^. Our analysis identified the restrictive residues of koala and mouse ACE2 in positions 31 and 353, respectively. Of note, humanization of koala or mouse ACE2 at either of these residues allowed these orthologs to bind viral spike proteins and mediate viral entry. Similarly, our previous study found that humanization of ACE2 from New World monkeys at positions 41 and 42 also resulted in gain-of-function phenotype^20^. Thus, together, our studies establish positions 31 (Lys), 41/42 (Tyr/Gln) and 353 (Lys) as genetic determinants regulating the usage of ACE2 as a receptor for SARS-CoV-2 infection (**Fig 5**). These data suggest that non-susceptible species have evolved species-specific mechanisms to restrict viral entry, and it is important to understand such mechanisms to develop novel animal models for SARS-CoV-2 research.

**Figure 5.**
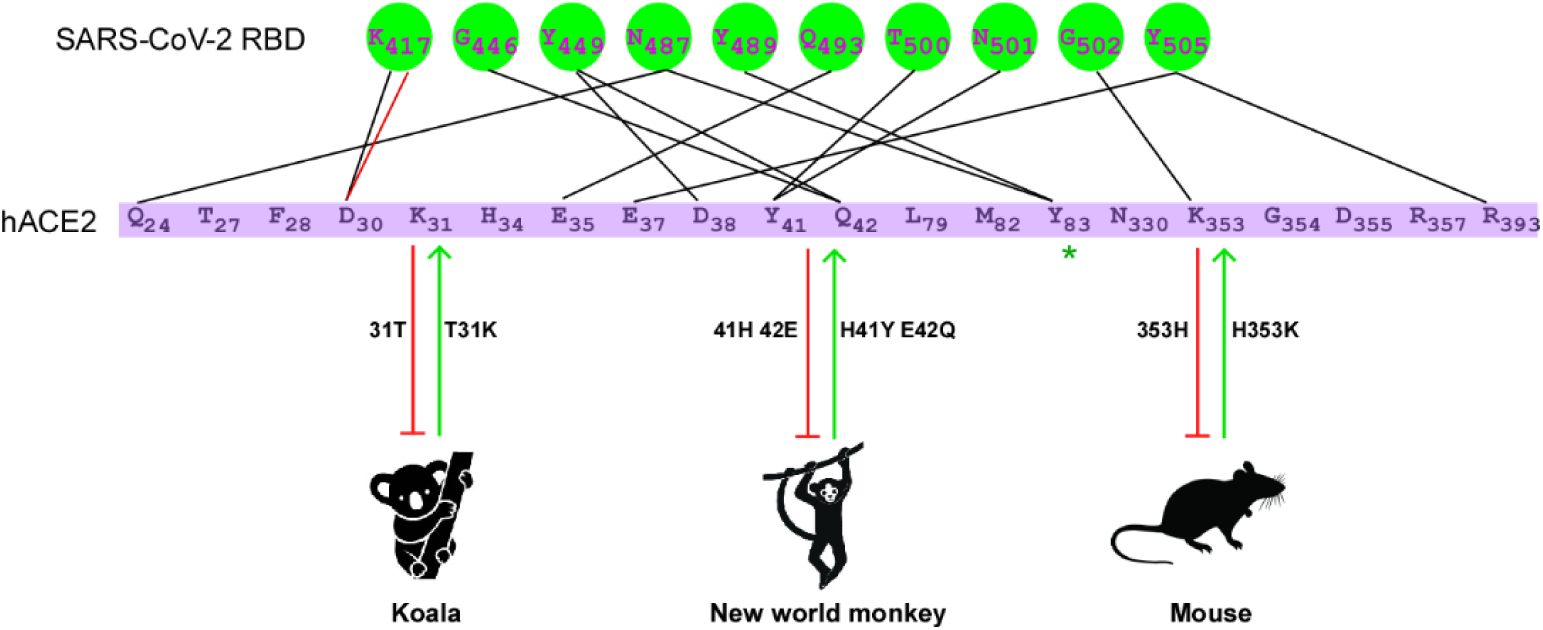
The genetic determinants of ACE2 required for SARS-CoV-2 entry. Mouse, koala, and New World monkey ACE2 cannot serve as functional receptors to support SARS-CoV-2 entry, as determined by different genetic restrictions. Position 31 in koala ACE2 is Thr whereas that in human is Lys. Substitution of Thr with Lys in koala ACE2 allowed for binding to the SARS-CoV-2 spike and viral entry. Different from koala ACE2, the genetic restriction of mouse ACE2 His353. Lys353 in human ACE2 can hydrogen bond with Gly502 of the SARS-CoV-2 spike protein, stabilizing the ACE2-spike complex. The presence of His at this position in mouse ACE2 disrupts this interaction. However, humanization of mouse ACE2 at position 353 renders the protein supportive of SARS-CoV-2 entry. The genetic determinants of New World monkey ACE2 were localized at positons 41 and 42 as we previously reported^20^. Thus, three genetic determinants of the ability of ACE2 to serve as the SARS-CoV-2 receptor were identified by comparative analysis of ACE2 orthologs, and the receptor activities of normally non-susceptible ACE2 orthologs could be rescued by genetic modification.

ACE2 is a major host factor determining the host range and tissue tropism of SARS-CoV-2 infection^13,14,20,26^. Our comparative analysis of the sequences and viral receptor activities of susceptible (human) or non-susceptible (koala and mouse) ACE2 orthologs sheds more light on understanding how ACE2 mediates SARS-CoV-2 entry and defines the species-specific genetic determinants of ACE2 governing viral receptor activity. Interestingly, we also compared the susceptibility of these orthologs and their humanized mutants to SARS-CoV. We found that mouse ACE2 bound SARS-CoV spike and mediated SARS-CoV pseudovirus entry, albeit with lower efficiency than human ACE2 (**Fig 2b and 3c**), but it did not bind SARS-CoV-2 spike (**Fig 2b and 3a**). Notably, koala ACE2 could not bind SARS-CoV or SARS-CoV-2 spike. Humanizing koala ACE2 rendered the protein only able to support SARS-CoV-2, but not SARS-CoV, entry as evidenced by our cell-based binding and pseudoviron entry experiments. Our findings highlight differential mechanisms of utilization of ACE2 as a receptor by SARS-CoV and SARS-CoV-2, which may contribute to the apparent ease of transmission and high mortalities of SARS-CoV-2.

There are still no prophylactic vaccines or effective antiviral drugs available to prevent SARS-CoV-2 infection and associated disease^35,36^. An animal model is needed to evaluate the efficacy of vaccine candidates and antiviral strategies against SARS-CoV-2 *in vivo*. Mice (*Mus musculus*) have long served as models of human biology and disease^37,38^. However, development of mouse models for COVID-19 has been challenging due to their natural resistance to SARS-CoV-2 infection^13,39^. Although human ACE2 transgenic mice can be infected with SARS-CoV-2, the non-physiological expression and tissue distribution of human ACE2 in such models represent a major limitation for conducting pathogenesis studies and accurately modeling the disease features observed in severe cases of COVID-19^16,17^. Thus, animal models which recapitulate human disease are urgently required. Genetic modifications to minimally humanize the hepatitis C virus (HCV) receptors CD81 and OCLN in mice by knocking in critical domains rendered the mice susceptible to HCV entry^40-42^. Based on our findings here, a similar knock-in strategy to generate an ACE2 humanized mouse is worth considering to aid in the development of novel mouse models capable of supporting SARS-CoV-2 infection as well as antiviral therapies.

Altogether, our study sheds more light on the species-specificity of interactions between SARS-CoV-2 and its cellular receptor ACE2 and uncovers evidence for a molecular arms race between SARS-CoV-2 and animal species. Here we identified the genetic basis defining the host range of SARS-CoV-2, which could potentially inform the development of novel animal models for SARS-CoV-2 research.

## Acknowledgements

We thank Di Qu, Xia Cai, Zhiping Sun, Wendong Han, and other colleagues at the Biosafety Level 3 Laboratory of Fudan University for help with experiment design and technical assistance. We thank Dr. Jenna M. Gaska for suggestions and revision of the manuscript. We are grateful to other members of the Ding lab for critical discussions and comments on the manuscript.

This work was supported by Tsinghua-Peking University Center of Life Sciences (045-61020100120), National Natural Science Foundation of China (32041005), National Key Research and Development Program of China (2020YFA0707701), Tsinghua University Initiative Scientific Research Program (2019Z06QCX10), Beijing Advanced Innovation Center for Structure Biology (100300001), Start-up Foundation of Tsinghua University (53332101319), Project of Novel Coronavirus Research of Fudan University (to Y.X.), and Development Programs for COVID-19 of Shanghai Science and Technology Commission (20431900401). Research in the Douam Lab at Boston University is supported by a Boston University/NEIDL Startup fund.

## Materials and methods

### Cell culture and SARS-CoV-2 virus

HEK293T (American Tissue Culture Collection, ATCC, Manassas, VA, CRL-3216), Vero E6 (Cell Bank of the Chinese Academy of Sciences, Shanghai, China) and A549 (ATCC) cells were maintained in Dulbecco’s modified Eagle medium (DMEM) (Gibco, NY, USA) supplemented with 10% (vol/vol) fetal bovine serum (FBS), 10mM HEPES, 1mM sodium pyruvate, 1×non-essential amino acids, and 50 IU/ml penicillin/streptomycin in a humidified 5% (vol/vol) CO2 incubator at 37°C. Cells were tested routinely and found to be free of mycoplasma contamination. The SARS-CoV-2 strain nCoV-SH01 (GenBank accession no. MT121215) was isolated from a COVID-19 patient and propagated in Vero E6 cells for use. All experiments involving virus infections were performed in the biosafety level 3 facility of Fudan University following all regulations.

### Plasmids

The cDNAs encoding the ACE2 orthologs were synthesized by GenScript and cloned into the pLVX-IRES-zsGreen1 vector (Catalog No. 632187, Clontech Laboratories, Inc) with a C-terminal FLAG tag. ACE2 mutants were generated by Quikchange (Stratagene) site-directed mutagenesis. All constructs were verified by Sanger sequencing.

### Lentivirus production

Vesicular stomatitis virus G protein (VSV-G) pseudotyped lentiviruses expressing ACE2 orthologs tagged with FLAG at the C-terminus were produced by transient co-transfection of the third-generation packaging plasmids pMD2G (Addgene #12259) and psPAX2 (Addgene #12260) and the transfer vector with VigoFect DNA transfection reagent (Vigorous) into HEK293T cells. The medium was changed 12 h post transfection. Supernatants were collected at 24 and 48h after transfection, pooled, passed through a 0.45-µm filter, and frozen at -80°C.

### Surface ACE2 binding assay

A549 cells were transduced with lentiviruses expressing the ACE2 of different species for 48 h. The cells were collected with TrypLE (Thermo #12605010) and washed twice with cold PBS. Live cells were incubated with the recombinant protein, the S1 domain of the SARS-CoV-2 spike C-terminally fused with Fc (Sino Biological #40591-V02H, 1μg/ml), at 4 °C for 30 min. After washing, cells were stained with goat anti-human IgG (H + L) conjugated with Alexa Fluor 647 (Thermo #A21445, 2 μg/ml) for 30 min at 4 °C. Cells were then washed twice and subjected to flow cytometry analysis (Thermo, Attune(^TM^) NxT).

### Production of SARS-CoV-2 or SARS-CoV pseudotyped virus

Pseudoviruses were produced in HEK293T cells by co-transfecting the retroviral vector pTG-MLV-Fluc, pTG-MLV-Gag-pol, and pcDNA3.1 expressing the SARS-CoV spike gene, SARS-CoV-2 spike gene or VSV-G (pMD2.G (Addgene #12259)) using VigoFect (Vigorous Biotechnology). At 48 h post transfection, the cell culture medium was collected for centrifugation at 3500 rpm for 10 min, and then the supernatant was subsequently aliquoted and stored at -80°C for further use. Virus entry was assessed by transduction of pseudoviruses in cells expressing ACE2 ortholog or mutants in 48-well plates. After 48 h, intracellular luciferase activity was determined using the Luciferase Assay System (Promega, Cat. #E1500) according to the manufacturer’s instructions. Luminescence was recorded on a GloMax® Discover System (Promega).

### Analysis of SARS-CoV-2 infection by high-content imaging system

A549 cells were transduced with lentiviruses expressing the ACE2 of different species for 48 h. Cells were then infected with nCoV-SH01 (SARS-CoV-2) at an MOI of 1 for 1 h, washed three times with PBS, and incubated in 2% FBS culture medium for 48 h. Cell were then fixed for viral antigen staining with 4% paraformaldehyde in PBS, permeablized with 0.2% Triton X-100, and incubated with a rabbit polyclonal antibody against the SARS-CoV nucleocapsid protein (Rockland, 200-401-A50, 1μg/ml) and a mouse anti-FLAG M2 antibody (Sigma-Aldrich #1804, 1μg/ml) at 4 °C overnight. After three washes, cells were incubated with a secondary goat anti-rabbit antibody conjugated with Alexa Fluor 555 (Thermo #A32732, 2 μg/ml) and goat anti-mouse antibody conjugated with Alexa Fluor 647 (Thermo #A21235, 2 μg/ml) for 2 h at room temperature, followed by staining with 4’,6-diamidino-2-phenylindole (DAPI). Images were collected using an Operetta High-Content Imaging System (PerkinElmer). For high-content imaging, three biological replicates for each ACE2 ortholog/variant on different 96-well plates were scanned, and five representative fields were selected for each well. Image analysis was performed using the PerkinElmer Harmony high-content analysis software 4.9. ACE2-transduced cells were automatically identified by DAPI (nuclei) and zsGreen (cytoplasm, the lentiviral vector containing the ACE2 orthologs also expressed zsGreen). The mean fluorescent intensity (MFI) of the Alexa 647 (ACE2 orthologs) and Alexa 555 (viral nucleocapsid) channels were subsequently calculated for each cell. Cells with an Alexa 647 channel MFI >1000 were gated as ACE2-positive cells. ACE2-positive cells with an Alexa 555 channel MIF >1600 were gated as SARS-CoV-2-infected cells. The formula of SARS-CoV-2-infected cells/ACE2-positive cells was followed as the readout for infection efficiency.

### Statistical analysis

One-way analysis of variance (ANOVA) with Tukey’s honestly significant difference (HSD) test was used to test for statistical significance of differences between the different group parameters. *P* values of less than 0.05 were considered statistically significant.

## Supplemental Figures

**Supplemental Figure 1.**
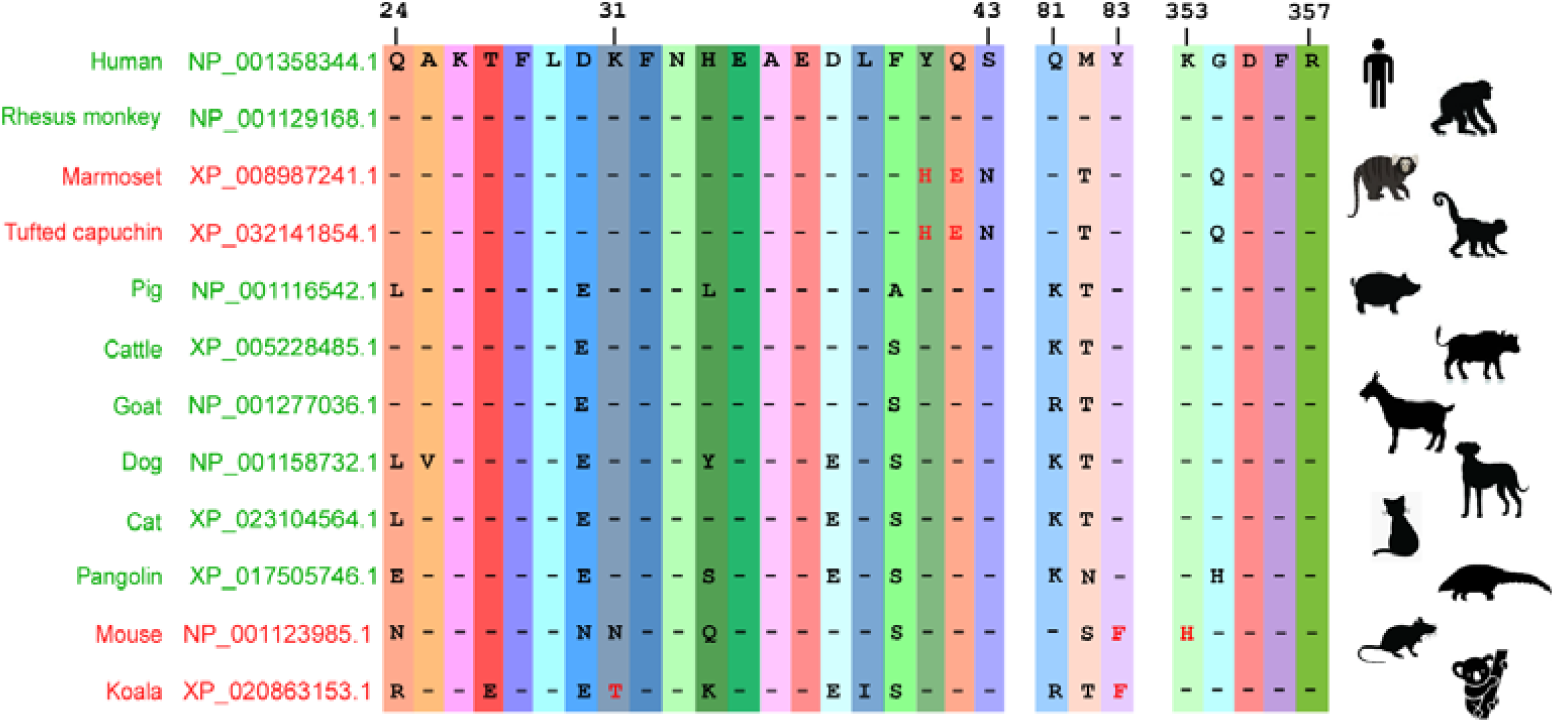

**Supplemental Figure 2.**
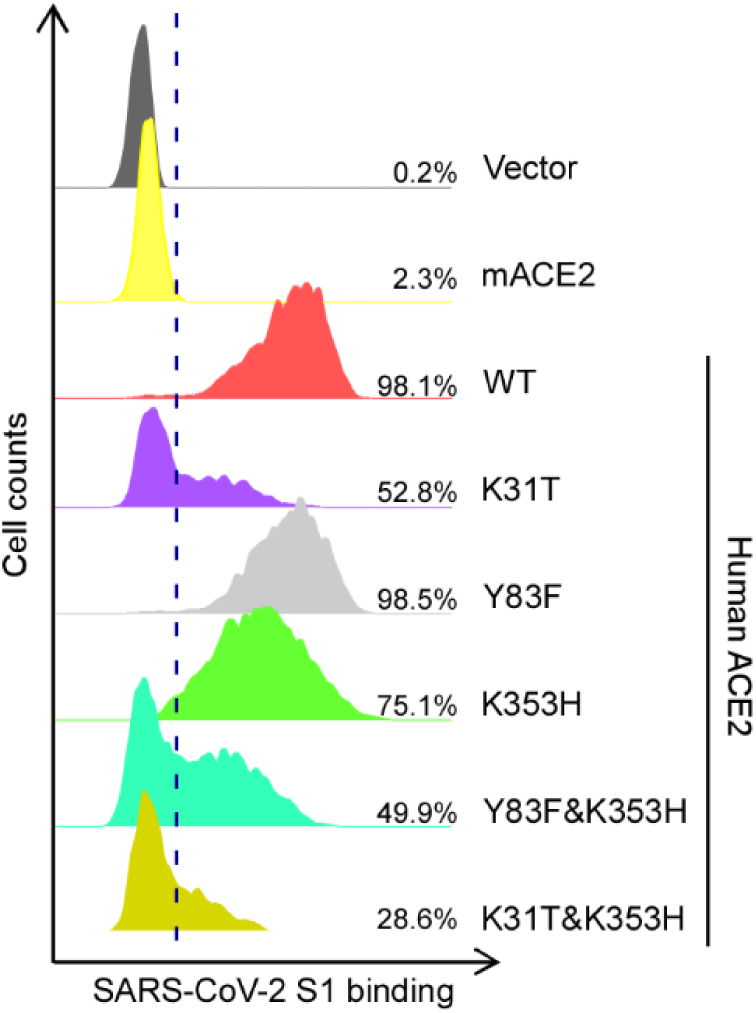

## Notes

### Competing Interest Statement

The authors have declared no competing interest.

